# RGS2 is an innate immune checkpoint for TLR4 and Gαq-mediated IFNγ generation and lung injury

**DOI:** 10.1101/2023.09.22.559016

**Authors:** Jagdish Chandra Joshi, Bhagwati Joshi, Cuiping Zhang, Somenath Banerjee, Vigneshwaran Vellingiri, Vijay Avin Balaji Raghunathrao, Lianghui Zhang, Ruhul Amin, Yuanlin Song, Dolly Mehta

## Abstract

IFNγ, a type II interferon secreted by immune cells, augments tissue responses to injury following pathogenic infections leading to lethal acute lung injury (ALI). Alveolar macrophages (AM) abundantly express Toll-like receptor-4 and represent the primary cell type of the innate immune system in the lungs. A fundamental question remains whether AM generation of IFNg leads to uncontrolled innate response and perpetuated lung injury. LPS induced a sustained increase in IFNg levels and unresolvable inflammatory lung injury in the mice lacking RGS2 but not in RGS2 null chimeric mice receiving WT bone marrow or receiving the RGS2 gene in AM. Thus, indicating RGS2 serves as a gatekeeper of IFNg levels in AM and thereby lung’s innate immune response. RGS2 functioned by forming a complex with TLR4 shielding Gaq from inducing IFNg generation and AM inflammatory signaling. Thus, inhibition of Gaq blocked IFNg generation and subverted AM transcriptome from being inflammatory to reparative type in RGS2 null mice, resolving lung injury.

**Highlights:**

- RGS2 levels are inversely correlated with IFNγ in ARDS patient’s AM.
- RGS2 in alveolar macrophages regulate the inflammatory lung injury.
- During pathogenic insult RGS2 functioned by forming a complex with TLR4 shielding Gαq from inducing IFNγ generation and AM inflammatory signaling.

**eToc Blurb:** Authors demonstrate an essential role of RGS2 in macrophages in airspace to promoting anti-inflammatory function of alveolar macrophages in lung injury. The authors provided new insight into the dynamic control of innate immune response by Gαq and RGS2 axis to prevent ALI.

## Introduction

Alveolar macrophages (AM) are the predominant tissue-resident macrophages in the lungs that become activated upon sensing pathogens in most lung disorders, including life-threatening diseases such as acute lung injury, ARDS, and COVID-ARDS (Bain and MacDonald, 2022; Dang et al., 2022; Epelman et al., 2014; Grant et al., 2021; Wendisch et al., 2021). Uncontrolled generation of IFNγ, a type II interferon, is consistently observed in the lungs of severe ALI/ARDS patients (Aoyagi et al., 2011; Verma et al., 2021; Yamada et al., 2004). However, the lymphocytes are considered primary producers of IFNγ in the context of anti-viral immunity (Choi et al., 2018; de Bruin et al., 2014; Lee and Ashkar, 2018; Varma et al., 2002).

Lipopolysaccharide, LPS, a cell wall component of all gram-negative bacteria, activates AM innate response through pattern recognition receptors such as toll-like receptors 4 (TLR4) and co-receptor CD14 (Joshi et al., 2020; Lu et al., 2015; Rayees et al., 2022; Rayees et al., 2020). Combining LPS with IFNγ boosted the massive release of proinflammatory and cytotoxic factors such as TNFα, IL6, and nitric oxide (NO) (Saleh et al., 2021). These synergistic effects of LPS and IFNγ were also observed *in vivo*, causing impaired injury resolution, which is also associated with ALI mortality (Dolowschiak et al., 2016; Gadotti et al., 2020). Moreover, a high amount of IFN-γ in severe SARS-CoV-2 patients is associated with PANoptosis and increased susceptibility to infection (Akamatsu et al., 2021; Heuberger et al., 2021). Thus, whether AM generation of IFNγ leads to uncontrolled innate response and perpetuated lung injury is widely unknown.

AM also express several G-protein coupled receptors that cooperate with LPS-TLR4 signaling to modulate AM activation and inflammatory responses via regulating Ca^2+^ and cAMP levels (Dauphinee et al., 2011; Fan et al., 2005; Li et al., 2015; Rayees et al., 2019; Solomon et al., 1998). The regulator of G-protein signaling (RGS) serves as an endogenous inhibitor of G proteins through its GTPase activating proteins (GAPs) action (Dong et al., 2017; Kehrl and Sinnarajah, 2002; McDonough et al., 2021; Osei-Owusu and Blumer, 2015; Philip et al., 2010). A prime example is RGS2, which can inhibit signaling mediated by Gαq and, to a lesser extent, Gαi and Gs protein (Ji et al., 2011; Klepac et al., 2019; Takimoto et al., 2009).

Here, we reveal an unexpected role of RGS2 as a critical determinant of IFNγ generation and innate immune response and establish its relevance in the murine model of ALI. We show that RGS2 levels are inversely correlated with IFNγ in AMs from ARDS patients and a preclinical mouse model of acute lung injury (ALI). RGS2 suppressed IFNγ levels and AM inflammatory signaling by inhibiting the interaction of TLR4 with Gαq. Thus, inhibition of Gαq blocked IFNγ generation and subverted AM transcriptome from being inflammatory to reparative type in RGS2 null mice, resolving lung injury. Our findings offer a novel perspective on the G-protein signaling in macrophages during lung injury, where RGS2 fine-tunes IFNγ generation and innate immune response in macrophages to resolve ALI.

## Results

### Loss of RGS2 leads to unresolvable inflammatory lung injury

To address the basis of RGS2 regulation of innate response, we first assessed publicly available data (GSE120000) and found that RGS2 was highly expressed in AM (**Fig. S1A-B**) (Mould et al., 2019; Tabula Muris et al., 2018). Further analysis showed that RGS2 expression decreased in AM at the peak of injury, associated with increased NOS2 expression (also known iNOS) (**Fig. S1B-C**), while enriched in reparative AM and airspace recruited macrophages during resolution of injury (**Fig. S1D**). Thus, using the endotoxemia model of lung injury in a WT mouse (0.5 mg/kg *i.t* LPS) (Dagvadorj et al., 2015), (**Fig. 1A**), we found that RGS2 mRNA expression in the AM and lungs were decreased at the peak of lung edema formation at 16h after LPS challenge and gradually returned to basal levels over 64h during the repair of injury (**Fig. 1B-D**). Furthermore, a similar decrease of RGS2 expression in the lungs was observed in COVID-19 mice models of lung injury (**Fig. 1E**).

**Figure 1.**
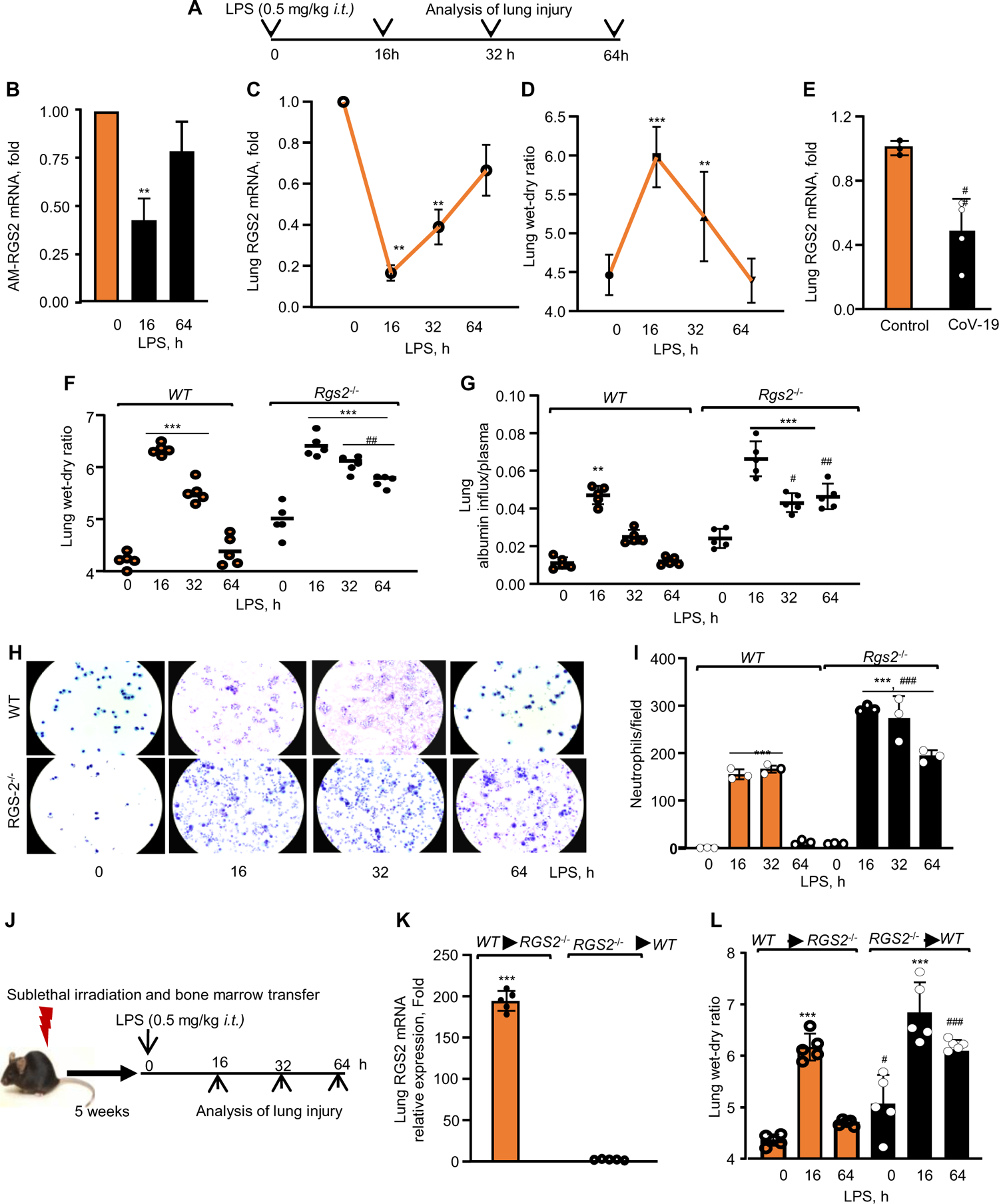
RGS2 is required for resolving lung inflammatory injury. **(A)** Schematics of lung injury determination after administration of LPS (0.5 mg/kg) *i.t.*, in control or RGS2 null mice. RGS2 expression in AM (**B**) and lungs (**C**) and lung edema (**D**) at indicated time points from LPS-exposed control mice. RGS2 mRNA was determined using qPCR and GAPDH as an internal control. Plots B-C show mean ± SD; n=3 mice/group. (**D**) Lung edema (n=5) was determined by measuring the wet-dry ratio. **(E)** SARS-Cov-2 infected Ace-2 and control mice lung cDNA was used to assess the RGS2 expression using qPCR (n=4 mice/group). Lung edema (**F**) and albumin influx in lung parenchyma versus plasma (**G**). Individual values along with mean ± SD are shown (n=5 mice/group). (**H-I)** Representative images of BALF stained with H&E is shown in H while plot shows individual values along with mean ± SD, (n=3 mice/group) of neutrophils/field. (**J)** WT mice were injected with RGS2 null bone marrow cells after irradiation and vice versa. Five weeks after the transplantation, the mice were challenged with *i.t.* LPS for indicated time points. (**K)** mean ± SD of RGS2 mRNA quantified as in B (n=5 mice/group). **(L)** Wet-dry ratio in the chimeric mice. The plot shows individual values with mean ± SD, n = 5 mice/group. ***p < 0.001 indicates significance relative to a control group receiving the vehicle alone. ##p < 0.01 indicates significance relative to corresponding mice post-LPS challenge. Analysis was performed using one-way ANOVA followed by Tukey’s multiple comparisons test.

We next subjected RGS2 null mice to the above endotoxemia model and observed that LPS similarly increased lung edema and vascular permeability in WT and RGS2 null mice at 16h (**Fig. 1F-G**). However, whereas WT mice resolved lung injury within the next 64 h, RGS2-lacking mice failed to do so as the lung edema and vascular permeability remained markedly higher (**Fig. 1F-G**). Also, RGS2 null lungs showed increased neutrophil influx over the time course of injury (**Fig. 1H-I**).

To determine the contribution of RGS2 expression in hematopoietic cells versus lung parenchyma in suppressing inflammatory injury, we lethally irradiated the mice and transplanted WT bone marrow cells to RGS2 null mice (WT→RGS2 chimera) or RGS2 null bone marrow into WT mice (RGS2→WT chimera) (**Fig. 1J**). The successful creation of radiation chimeras was confirmed by quantifying RGS2 expression at RNA and protein levels in chimeric mice lungs five weeks after bone marrow transplantation (**Fig. 1J-K and Fig. S1E)**. We then administered LPS to these chimeric mice and found that WT → RGS2 null chimeric mice resolved injury while RGS2 null →WT chimeric mice did not (**Fig. 1L**). These results suggest that the absence of RGS2 in nonhematopoietic host tissue was not critical to resolving lung injury, whereas RGS2 deficiency in the recipient hematopoietic cells was sufficient for inducing long-lasting lung injury.

### RGS2 negatively regulates IFNγ generation in the lung macrophages

The inverse correlation between RGS2 expression and NOS2 prompted us to perform RNA-seq analysis in lavage macrophages to assess the transcriptional machinery responsible for programming RGS2 null macrophages into the inflammatory lineage. We focused on assessing transcriptome at 64h post-LPS challenge and found that around 1200 genes were upregulated in WT or RGS2 null macrophages than unchallenged macrophages. Gene ontology (GO) biological process (BP) showed the immune system process and innate immune response as the top-listed pathways in activated WT or RGS2 null macrophages (**Fig. 2A-B** and **Fig. S2A-B**). We screened the top 25 upregulated genes and found increased expressions of IFNGR1, NOS2, and guanylate-binding proteins (Gbp4, Gbp5, and Gbp9) in RGS2 null AM than in WT-AM (**Fig. 2A-B**). We also found an increase in the expression of transcription factor IRF7, which belongs to the member of the interferon regulatory transcription factor family (IRF) (Farlik et al., 2012; Ning et al., 2011) in the RGS2 null AM than WT-AM (**Fig. 2A-B**). Consistently, a 2-fold increase in IFNγ mRNA was observed in RGS2 null AM basally than WT-AM (**Fig. 2C**). RGS2 deletion caused a leftward shift in the time course of IFNγ generation by LPS in AM leading to a massive increase at 64h (**Fig. 2D**). The levels of IFNγ were also increased 9-fold in SARS-CoV-2 infected lungs (**Fig. 2E**). In other studies, we administered LPS in LyzM-GFP mice, where all myeloid cells are green to determine AM-RGS2 and IFN-g at protein levels using confocal analysis. These data also demonstrated that LPS significantly increased IFNγ levels while concomitantly decreasing RGS2 protein (**Fig. 2F-H**).

**Figure 2.**
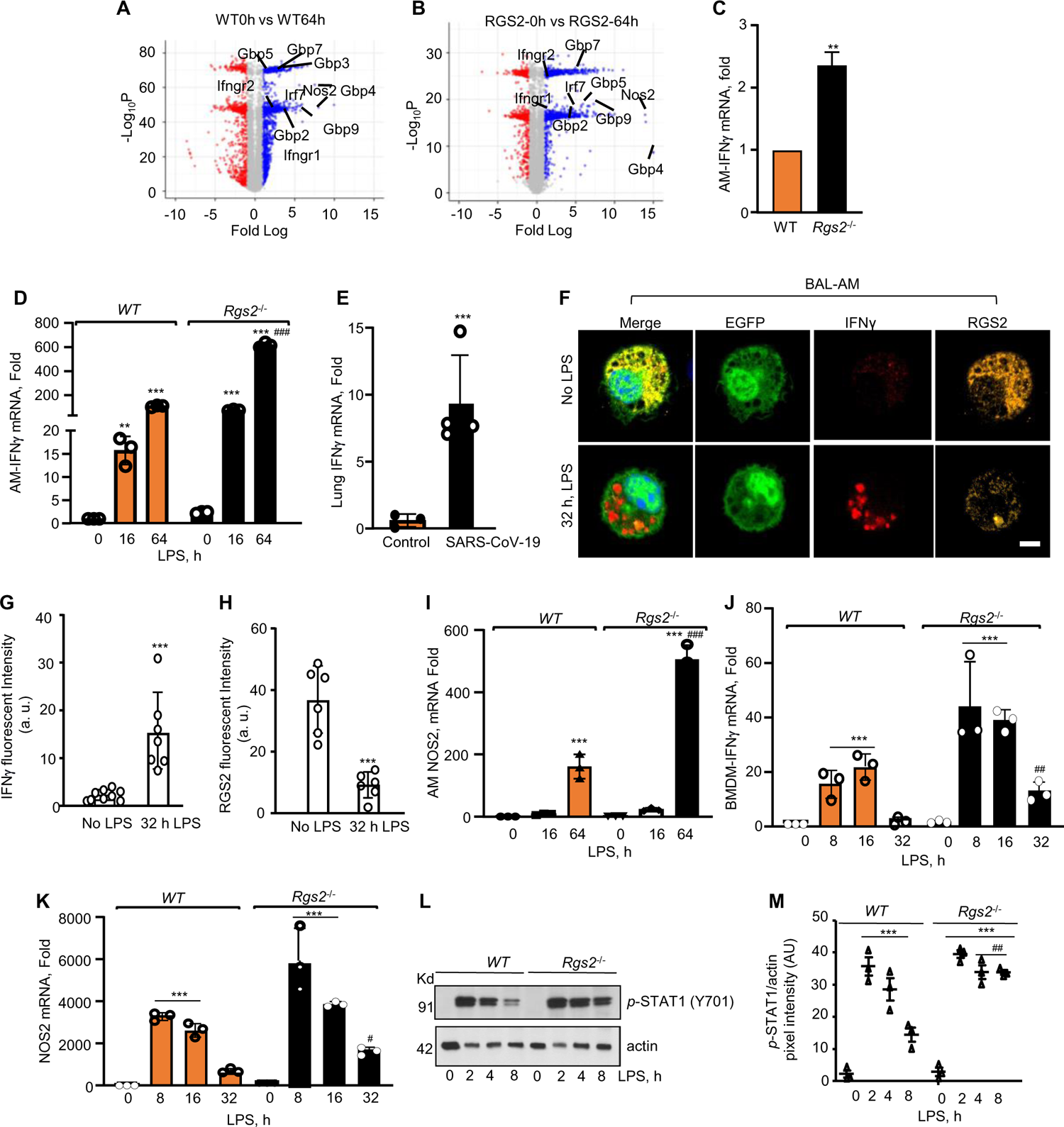
Loss of RGS2 in macrophages impairs lung inflammatory injury by generating IFNγ. (**A-B**) Volcano plot of AM transcriptome in WT or RGS2 null AM after without or with LPS challenge (n=3 mice/group). **(C-E)** IFNγ expression in AM at indicated times post LPS administration in mice (C-D) or in lungs of mice 7 days after receiving SARS-Cov2 (E). Using qPCR, expression was determined with GAPDH as an internal control (n=3 mice/group). **(F-H)** AM isolated from Lyz-M mice 32 h after administration of LPS were stained with anti-RGS2 or anti-IFNγ antibodies. Images were acquired using confocal microscopy (n=3 mice/group). Representative images are shown in panel F while plots show the quantitation of fluorescent intensity of IFNγ (**G**) and RGS2 (**H**) in AM. **I,** NOS2 expression in AM using qPCR (n=3 mice/group). GAPDH was used as an internal control. **(J-K)** WT and RGS2 null BMDM were stimulated with LPS (1μg/ml) for indicated time points, and expression of IFNγ (**J**) and NOS2 (**K**) expression were measured using qPCR (n=3 mice/group) as in Figure 1A-B. **(L-I)** Phosphorylation of STAT1 (Y701) after without or with LPS stimulation of BMDM. A representative immunoblot is in panel L, while densitometry analysis is shown in panel M (n=3 experiments/group). Actin was used as a loading control. Plot C shows mean ± SD, while Data in D-E, G-K, and M shows individual values with mean ± SD, n = 5 mice/group. **p < 0.01, ***p < 0.001 indicates significance relative to a control group receiving vehicle alone. #p <0.05 and ##p <0.01 indicates significance relative to corresponding mice post-LPS challenge. Analysis was performed using one-way ANOVA followed by Tukey’s multiple comparisons test.

IFNγ binds its receptors which then recruit the tyrosine kinase JAK2. JAK2 activates STAT proteins to induce NOS2 (Fenton et al., 1997; Fukao et al., 2001; Rothfuchs et al., 2001). We found that LPS induced a 5-fold higher expression of NOS2 in RGS2-deficient AM than the WT-AM (**Fig. 2I**). These changes were also apparent in bone marrow-derived macrophages (BMDM) from RGS2 null mice versus WT-BMDM. LPS increased IFNγ and NOS2 in WT-BMDM at 8 h, which increased further at 16 h but then returned to basal level at 32h (**Fig. 2J-K**). However, compared to WT-BMDM, LPS markedly augmented the increase of IFNγ and NOS2 in RGS2 null BMDM, which remained higher even at 32h (**Fig. 2J-K**). LPS increased the phosphorylation of STAT1 at two h, which returned to near basal level by 8h in WT-BMDM (**Fig. 2L-M**). However, the phosphorylation of STAT1 was persistently increased in RGS2 null BMDM (**Fig. 2L-M**). LPS did not affect the levels of STAT2, STAT3, STAT5, and STAT6 phosphorylation **(Fig. S2C)**.

To evaluate whether RGS2 and IFNγ production is clinically relevant, we measured IFNγ in the BAL fluid of ARDS patients in the ICU versus non-ARDS patients (referred to as controls) (**Fig. S3A**). ARDS patients exhibited more augmented IFNγ levels in their BAL fluids than controls (**Fig. 3A**). We also isolated BAL macrophages from ARDS and control patients and compared the relationship between RGS2 and IFNγ. We found a significant correlation between IFNγ levels with that of RGS2 in control and severe ARDS patients. The increase in RGS2 expression was correlated with a decrease in IFNγ levels in these patients (**Fig. 3B**). This inverse relationship was not observed in control patients (**Fig. 3B**).

**Figure 3.**
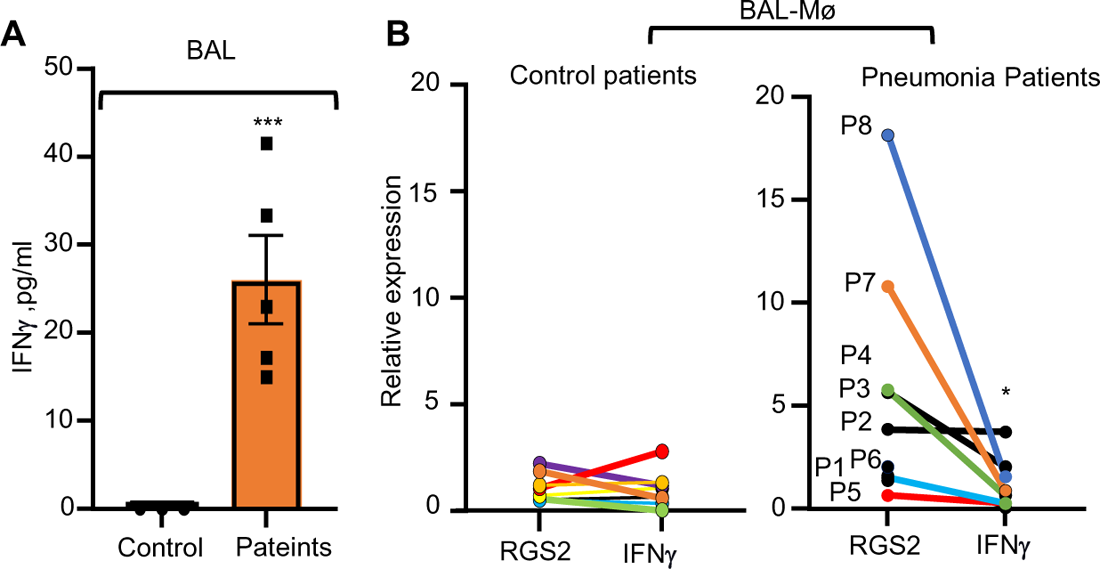
Lung injury alters RGS2 and IFNγ expression in ARDS AM. **(A)**, BAL-IFN**γ** levels and RGS2/IFN**γ** mRNA (**B**) in AM of control and ARDS patients. The data in plots A-B, show mean ± SD. *p < 0.05, ***p < 0.001 indicates significance relative to a control group. Analysis was performed using one-way ANOVA followed by Tukey’s multiple comparisons test.

### Rescuing RGS2 expression in macrophages resolves inflammatory lung injury in RGS2 null mice

To corroborate the causal role of RGS2 in macrophages in dampening IFNγ levels and inflammation, we complexed RGS2 cDNA with liposomes and delivered these lipid nanoparticles *i.v.* into RGS2 null mice, as described in **Fig. 4A**. We chose *i.v.* route because RGS2 is highly expressed in reparative AM and airspace-recruited macrophages during resolution of injury (Mould et al., 2019; Tabula Muris et al., 2018). Using lung lysates and BAL macrophages, we confirmed that RGS2 was delivered and expressed in the macrophages (**Fig. 4B-D**). Rescue of RGS2 expression in RGS2 null mice dampened IFNγ and NOS2 expression by LPS (**Fig. 4E-F**). However, this response was not seen in RGS2 null mice transducing vectors only (**Fig. 4E-F**). Similarly, we observed that RGS2 null BMDM expressing RGS2 cDNA showed markedly less IFNγ generation than RGS2 null BMDM expressing vector alone (**Fig. 4G-I**). Notably, RGS2 null mice receiving RGS2 cDNA but not vector alone resolved lung edema (**Fig. 4J**). These findings indicate that RGS2 expression plays a crucial role in controlling innate immune response and promoting injury resolution.

**Figure 4:**
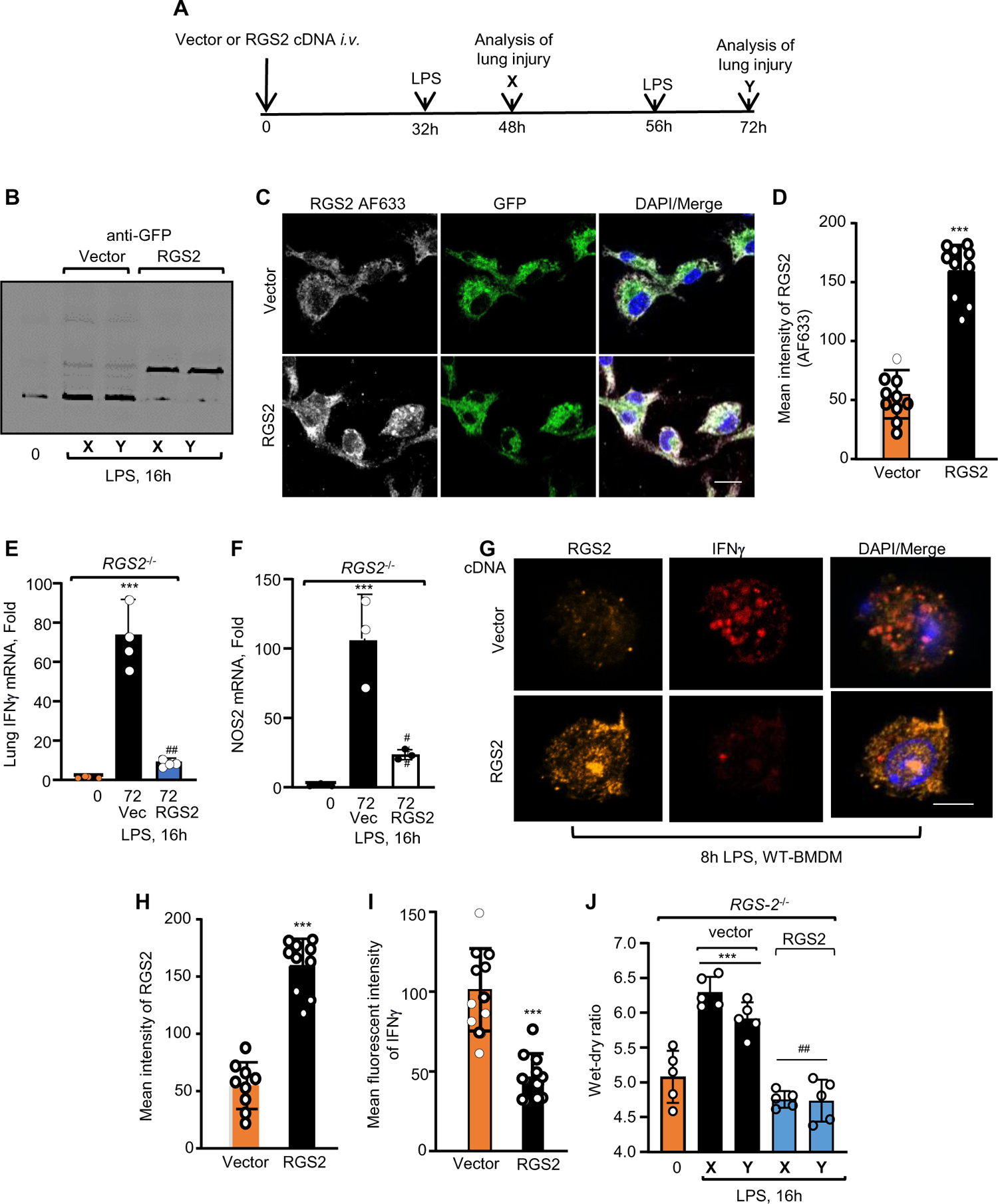
Restoration of RGS2 expression in lung AM resolves lung injury. **(A)** Schematics of RGS2 gene delivery in RGS2 null mice to assess its effect on edema and gene expression after LPS challenge. (**B-C)** A representative immunoblot (B) or image (C) from multiple experiments shows RGS2 expression in lungs receiving vector or GFP-tagged RGS2 cDNA. Lung lysates in “B” were immunoblotted with an anti-GFP antibody to detect RGS2 expression. X=48 h after liposomes and LPS; Y=72 h liposomes and LPS. BAL fluid in “C” was isolated from RGS2 null mice following 48h of liposomes delivery, and adhered AM were stained for RGS2 antibody (n=3 mice/group). (**D)** Fluorescent intensity of RGS2 in indicated group. **(E-F)** Lung IFN**γ** and NOS2 expression in an indicated mouse (n=3 mice/group). GAPDH was used as an internal control. (**G-I): (G)** A Representative image and **(H-I)** fluorescent intensity of RGS2 and IFN**γ** expression in BMDM transducing vector and RGS2-cDNA post 8h LPS (1 μg/ml) stimulation. Experiments were repeated at least three times independently. **(J)** Plot shows individual values along with mean ± SD of wet-dry ratio in the lungs of mice transducing indicated cDNA post 16h LPS challenge. n=5 mice/group. ***p < 0.001 indicates significance relative to a control group receiving vehicle alone. ##p < 0.01 indicates significance relative to a corresponding mouse post-LPS challenge. Analysis was performed using one-way ANOVA followed by Tukey’s multiple comparisons test. The data in plots B and E-F, show mean ± SD. ***p < 0.001 indicates significance relative to control group receiving vehicle alone. ##p <0.05 and ###p <0.01 indicates significance relative to corresponding mice post LPS and vector challenge. Analysis was performed using one-way ANOVA followed by Tukey’s multiple comparisons test.

### RGS2 suppression of Gαq activity resolves lung inflammation

RGS2 primarily inhibits Gαq activity, the latter a crucial determinant of intracellular Ca^2+^ levels (Kehrl and Sinnarajah, 2002). Therefore, we measured intracellular Ca^2+^ levels in WT and RGS2 null BMDM using a Ca^2+^ sensor, gCaMP-X, following LPS stimulation. LPS increased Ca^2+^ levels in RGS2 null mice BMDM more than in WT-BMDM **(Fig. S4A-B)**. Next, we tested the hypothesis that inhibiting Gαq activity in RGS2 null mice would rescue innate immune response and resolve lung injury by subverting the macrophage transcriptome from inflammatory to reparative phenotype. We used BIM-46187, which explicitly blocks Gαq activity (Schmitz et al., 2014) and confirmed that inhibition of the Gαq prevented the rise in cytosolic Ca^2^(McDonough et al., 2021; Philip et al., 2010) **(Fig. S4C).** Notably, BIM-46187 steeply reduced IFNγ generation in RGS2 null BMDM to the level seen in WT-BMDM (**Fig. 5A**). As Gαq activity subverted IFNγ level, we administered LPS in mice and 2h later, BIM-46187 was given *i.v.* (**Fig. 5B**). We observed that inhibition of Gαq reduced IFNγ generation in WT lungs by 40% and ∼80% in RGS2 null lungs (**Fig. 5C**). Consistently, BIM promoted resolution of edema formation even at the peak of injury in WT and RGS2 null mice (**Fig. 5D**). BIM alone had no effect on IFNγ generation or edema formation in WT or RGS2 null mice (data not shown). Transcriptomic and GO pathway analysis indicated that inhibition of Gαq targeted IFNγ cascade as IFNGR1, IFNGR2, IFNγ inducible genes (GBP-2, GBP-3, GBP-4, GBP-7, and GBP-9), NOS2, and IRF7 were reduced by BIM in RGS2 null macrophages (**Fig. 5E-5F**, Fig. S5A). We also transfected Gαq mutant (Q209L) devoid of binding RGS2 into the WT-BMDM and found that Gαq mutant transducing BMDM generated 3-fold higher IFNγ in response to LPS than BMDM expressing vector alone (**Fig. 5G**). The expression of Gαq transfection was confirmed by protein expression **(Fig. S5C).**

**Figure 5:**
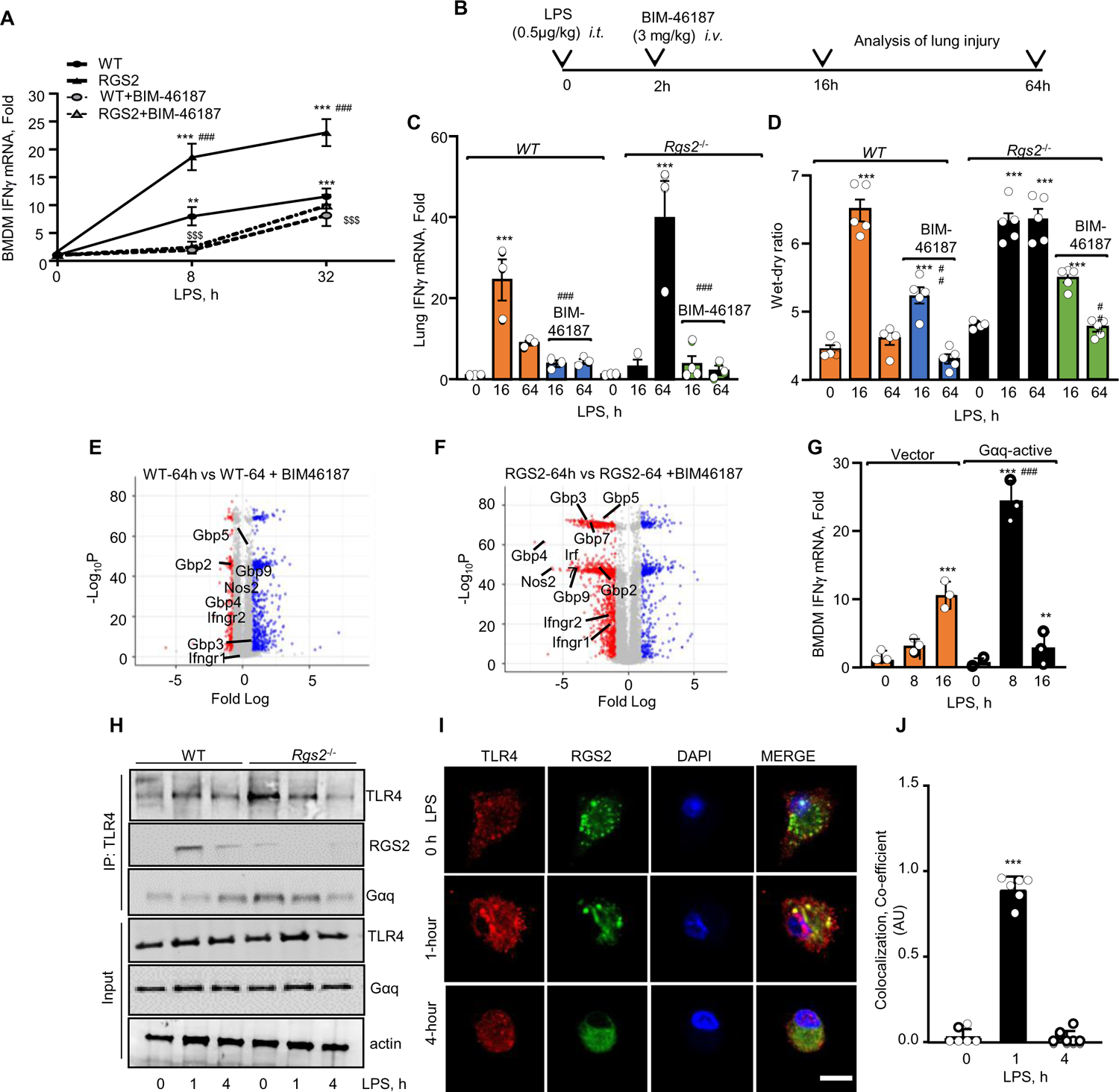
RGS2 resolves lung injury by limiting Gαq access to TLR4. **(A)** IFN**γ** expression in WT or RGS2 null BMDM after pretreatment with GαQ inhibitor BIM-46187 and LPS stimulation. BMDM were treated with BIM46187 (10 µg/ml) for 1h, after which IFN**γ** expression was determined using qPCR with GAPDH as an internal control (n=3 mice/group). **(B-D)** Schematics to assess the effects of BIM-46187 treatment on IFN**γ** expression (C) and LPS-induced lung injury in control and RGS2 null mice (D). Edema and gene expression was determined as described in Figures 1-2. (n=5 mice/group). **(E-F)** Volcano plot of WT and RGS2 null AM transcriptome after LPS and BIM-46187 treatment (n=3 mice/group). **(G)** IFN**γ** expression in WT-BMDM transducing vector and active Gαq post-LPS stimulation using qPCR (n=3 mice/group). GAPDH was used as an internal control. **(H)** Cell lysates from LPS-stimulated WT-BMDM were immunoprecipitated using TLR4 antibody, followed by immunoblotting with RGS2, TLR4, or Gαq to assess their interaction. A representative blot from two independent experiments is shown. **(I-J)** A representative image shows the co-localization of RGS2 with TLR4 in WT-BMDM stimulated with LPS (1µg/ml). Plot in I show co-localization coefficient of TLR4 and RGS2. Experiments were repeated three independent times.

LPS induces inflammatory signaling primarily by activating TLR4 (Joshi et al., 2021; Lu et al., 2015; Rochford et al., 2021). Because TLR4 signaling forms complex with Ca^2+^ dependent machinery (Park et al., 2016; Tauseef et al., 2012), we surmised that RGS2 limits Gαq access to TLR4 to regulate IFNγ synthesis. Hence, we used lysates from WT or RGS2 null BMDM post LPS stimulation and performed immunoprecipitation assay using TLR4 antibody. We found that in WT-BMDM, TLR4 interaction increased with RGS2 within 1 h after LPS stimulation decreasing at 4h (**Fig. 5H-J**). However, under these conditions, TLR4 weakly interacted with Gαq. Interestingly, TLR4 interacted with Gαq in RGS2 null BMDM basally which did not alter after LPS challenge (**Fig. 5H-J**). These findings indicate that RGS2 shields Gαq from TLR4 to timely control regulate IFNγ synthesis and innate immune response.

## Discussion

In the present study, we demonstrated the heretofore unknown role of RGS2 in safeguarding innate immune response in macrophages to resolve inflammatory lung injury. We showed that the loss of RGS2 programmed AM transcriptome towards inflammatory lineage due to upregulation of IFNγ cascade downstream of Gαq. Thus, mice lacking RGS2 failed to resolve lung injury, but inhibiting Gαq activity subverted AM transition to reparative macrophages, resolving lung injury. Therefore, our data demonstrate the fundamental innate immune-protective role of the RGS2 in AM to resolve ALI.

Fine-tuning of IFNγ generation in the lung milieu and downstream inflammatory signaling is a crucial determinant of ALI and other inflammatory diseases known (Akamatsu et al., 2021; Dolowschiak et al., 2016; Gadotti et al., 2020; Heuberger et al., 2021). Initial events that start in AM upon sensing pathogens is the burst of cytokines through the activation of TLR4, which recruits neutrophils and monocytes to defend the host. The airspace-recruited monocytes then turn into macrophages, which, together with AM repair the damaged lung, resolving lung injury (Bain and MacDonald, 2022; Dang et al., 2022; Jakubzick et al., 2017; Joshi et al., 2021; Lu et al., 2015; Wendisch et al., 2021). Gαq and RGS2 are highly expressed in AM at homeostasis and in reparative macrophages (Kleinjan et al., 2023; Rayees et al., 2019; Rayees et al., 2020). RGS2 holds Gαq inactive by inducing its GTPase activity (Ross and Wilkie, 2000). RGS2 can also inhibit Gαi and Gαs (Ji et al., 2011; Klepac et al., 2019). Using RGS2 null mice, we could show the influence of uncontrolled Gαq signaling in sustaining IFNγ generation, neutrophil accumulation, and lung innate immune response following endotoxemia. We also showed that preventing activation of Gαq promoted resolution of pulmonary edema in RGS2 null mice post LPS challenge implying that RGS2 functioned in AMs by blocking Gαq activity. Thus, we have identified RGS2 in AM as a suppressor of Gαq activity and resultant IFNγ generation and lung inflammatory injury. RGS2 is shown to be involved in regulating several organ functions, including blood pressure, airway reactivity, viral immunity, and kidney fibrosis (Dong et al., 2017; George et al., 2018; Jang et al., 2014; Kehrl and Sinnarajah, 2002; Klepac et al., 2019). We showed that expression of RGS2 decreased in AM during injury by LPS but restored to basal level during the resolution phase. In line with this, we showed that IFNγ-cascade was upregulated in AM lacking RGS2, and these mice failed to resolve lung injury showing that downregulation of the RGS2 in AM is a crucial mechanism underlying LPS-induced inflammatory injury. The mechanisms reducing RGS2 and leading to the emergence of IFNγ-linked transcriptome in AM following endotoxemia need to be clarified. A possibility is that epigenetic mechanisms such as DNA methylation are involved in reducing RGS2 synthesis in AM, as in the LPS injury model (Bouvet et al., 2018; Tuggle et al., 2014).

Gαq increases cytosolic Ca^2+^ downstream of several receptors (Chow and Davis, 2000; Lee et al., 2007; Scott et al., 2003). Findings from the current study identified a crucial role of Gαq in augmenting IFNγ-linked cascade in macrophages. So far, T cells are considered the primary source of IFNγ, which then polarizes macrophages to inflammatory lineage (Choi et al., 2018; de Bruin et al., 2014; Lee and Ashkar, 2018). Our data showed that WT-AM transiently but modestly generated IFNγ after the LPS challenge, associated with decreased RGS2 expression. IFNγ then restored to near basal level in association with resolving lung injury. However, loss of RGS2 massively induced IFNγ generation, impairing lung injury resolution. Thus, these studies imply that RGS2 controls IFNγ releases from the macrophages and resulting lung damage by repressing the Gαq signaling cascade to fine-tune the innate response. IFNγ activates JAK2-STAT signaling upon binding its receptors (Jones et al., 2020; Zhou et al., 2021). We also showed augmentation of STAT1 phosphorylation and NOS2 generation in RGS2 null macrophages showing IFNγ functioned in an autocrine/paracrine manner to induce inflammatory signaling in AM and lung injury. A caveat in delivering RGS2 gene or Gαq inhibitor, *i.v.,* is that these approaches also alter RGS2 or Gαq signaling in other cell types. However, we showed that overexpression of Gαq mutant lacking RGS2 binding site also induced IFNγ. Likewise, restoration of RGS2 or inhibition of Gαq reversed IFNγ levels in macrophages despite the LPS challenge.

An important question is how RGS2 controlled LPS-induced IFNγ generation in macrophages. We previously showed that TLR4 effector, MyD88 complexes with MLCK in a Ca^2+^-dependent manner (Srivastava et al., 2020). Since RGS2 suppressed the rise in cytosolic Ca^2+^, we surmised RGS2 binds TLR4 to limit Gαq activation by LPS. We showed TLR4 interacted with RGS2 but not Gαq following LPS stimulation. However, TLR4 interaction with Gαq increased in RGS2 null BMDM. Therefore, we interpreted that inefficient coupling between TLR4 and Gαq in macrophages enforced by RGS2 controlled IFNγ generation by LPS. Indeed, we showed that over-expressing constitutive active Gαq augmented IFNγ generation by LPS in WT macrophages, thus supporting the concept that RGS2 limits IFNγ generation by shielding Gαq from TLR4.

IFNγ regulates innate and autoimmunity and can lead to chronic tissue injury (Ivashkiv, 2018) (Lees, 2015). We observed increased IFNγ levels in the alveolar lavage of ARDS patients compared to controls, findings consistent with previous reports (McKeone et al., 2020; Paats et al., 2013). Interestingly, we found a reciprocal relationship between the expressions of RGS2 versus IFNγ in AM of ARDS. We showed that inhibition of Gαq induced the generation of reparative macrophages in RGS2 null mice and resolved injury even within 16h hours as opposed to the endogenous system requiring 64h, suggesting a means of enhancing kinetics of reparative AMs generation and the repair process.

In summary, findings from the current study provide novel insights into the mechanism that limits IFNγ generation in macrophages enabling timely resolution of inflammatory injury and lung repair. Targeting the IFNγ pathway in ARDS patients has yet to be successful. Therefore, selectively interfering with the RGS2-Gαq pathway may suppress IFNγ levels and promote lung injury resolution.

## SUPPLEMENT INFORMATION

Supplement information includes five figures.

**Supplementary Figure 1.** RGS2 expression in lung cells using publicly available data. **(A)** Data acquired from tabula muris data**. (B-C)** Violin plots show RGS2 and iNOS (Nos2) expression in lung AM and recruited macrophages under basal conditions and after LPS-induced injury and resolution. (**C**) UMAP of various macrophage populations of in BALF at the homeostasis, injury, and resolution phase. **(D)**, Immunoblot shows RGS2 protein expression in the lungs of chimeric mice after bone marrow transplantation mice. A representative blot is shown from experiments that were repeated two times.

**Supplementary Figure 2.** GO pathways analysis in macrophages lacking RGS2. **(A-B)** Shows upregulated genes in WT and RGS2 null AM after 64h of LPS exposure of mice (n=3 mice per group)**. (C)** Phosphorylation of STAT1 (Y701), STAT2, STAT3 (S727), STAT3 (Y705), STAT5 (Y694), and STAT6 (Y641) in indicated lung lysates was determined following without or with LPS stimulation of BMDM cells. A representative immunoblot is shown from experiments that were independently repeated three times. Actin was used as a loading control.

**Supplementary Figure 3.** Clinical state of ARDS patients in ICU admission. Lab and clinical diagnosis and characteristics of patients. Each patient’s lung CT scan is shown in the bottom.

**Supplementary Figure 4.** RGS2 blocks GαQ. **(A)** A representative image of cytosolic calcium from WT and RGS2 BMDM transfected with GFP tagged gCaMP-X following LPS stimulation at indicated time points (n=3 per set done independently). **(B)** Plot shows mean ± SD of changes in the fluorescence intensity of intracellular calcium. **(C)** Plot of individual data points along with mean ± SD of calcium increases in WT and RGS2 null BMDM following treatment with BIM and thrombin stimulation (n=6). **p < 0.01 indicates significance relative to the control group receiving the vehicle alone. Analysis was performed using one-way ANOVA followed by Tukey’s multiple comparisons test.

**Supplementary Figure 5.** Pathway analysis of Gαq-induced genes in AM following endotoxemia. **(A)** Downregulated AM pathway after BIM treatment of LPS-exposed mice (n=3 mice/group). **(B)** A representative blot of BMDM transducing vector or Gαq (n=3 mice/group).

## ACKNOWLEDGEMENTS

We thank Drs. Asrar B Malik, Department of Pharmacology, University of Illinois, for his feedback during the progression of the study, and Petr Borz, Department of Biochemistry, University of Lausanne, Switzerland, for providing GFP-tagged gCaMP-X calcium sensor. This work was supported by the NIH grants PO1-HL151327, PO1-HL160469, RO1-155941, RO1-HL084153, and RO1-HL165263. The authors have no conflicting financial interests.

## AUTHOR CONTRIBUTIONS

JCJ and DM conceptualized and designed the research. JCJ, BJ, SB, CZ, VV, VABR, LZ, and RA performed research. JCJ performed the mice work. BJ performed immunoprecipitation and western blotting. VV performed confocal imaging, SB and RA performed qPCR. VABR analyzed the PanSeq data, created the Volcano plots, and conducted pathway analysis. LZ performed the SARS-CoV-2 mice experiments. YS diagnosed and recruited ARDS patients. CZ collected the BALF samples from the hospital and processed them for experiments and analysis. JCJ and DM analyzed, interpreted, and wrote the manuscript.

## STAR* METHODS

### KEY RESOURCE TABLE

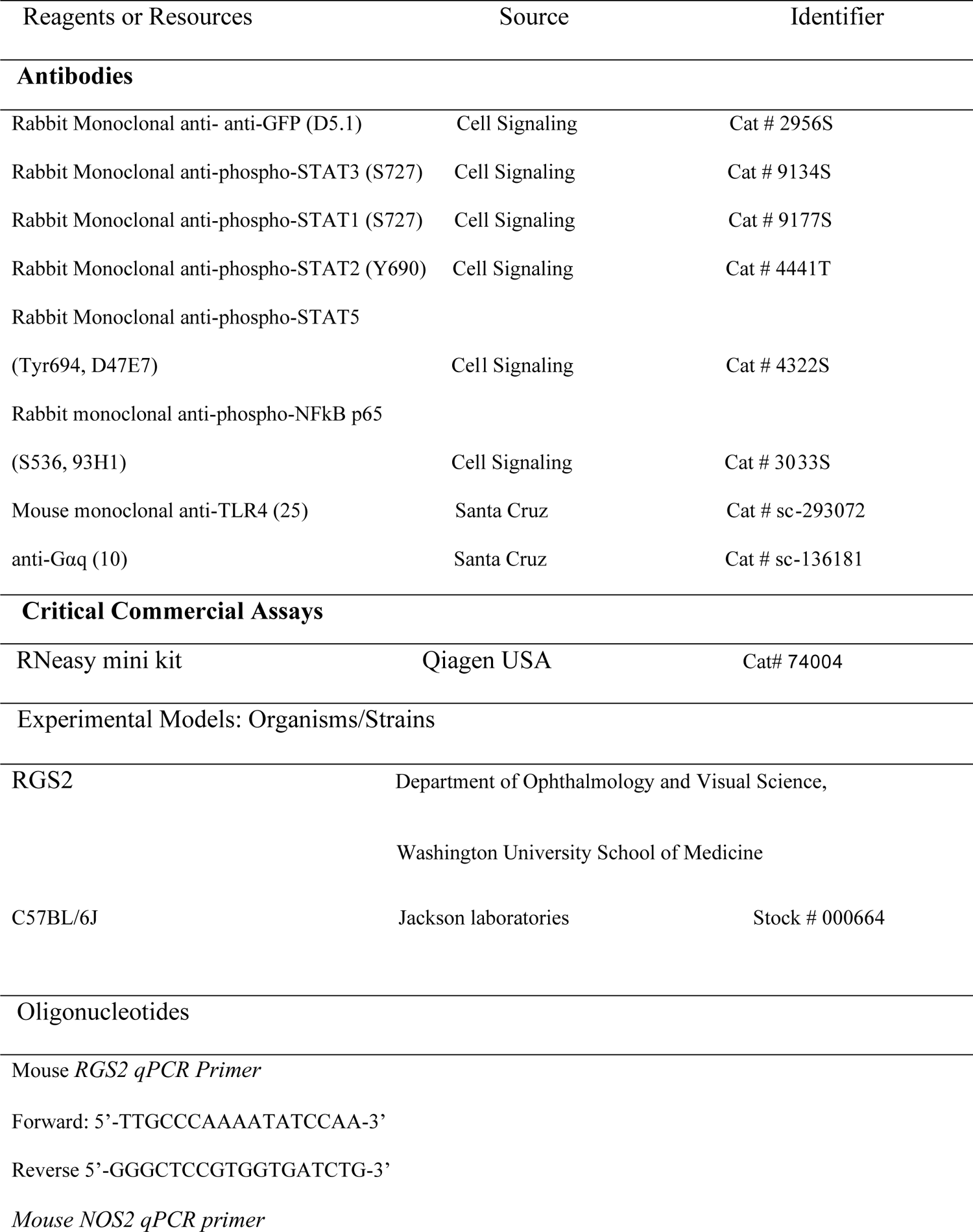
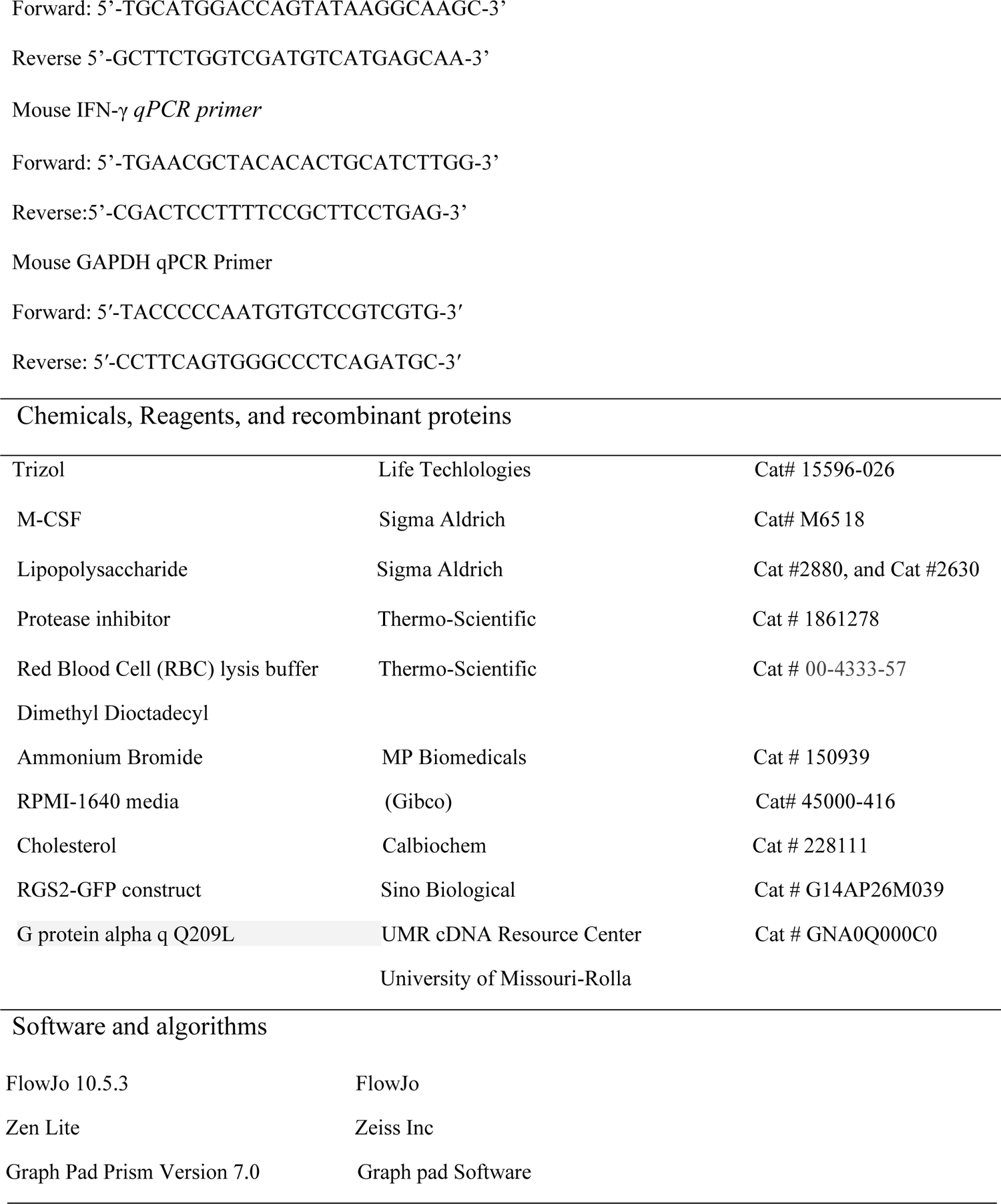

### CONTACTS FOR REAGENTS AND RESOURCE SHARING

Further information and requests for resources and reagents should be directed to and will be fulfilled by the Lead Contact, Dolly Mehta

### EXPERIMENTA MODEL

#### Animals

The animals used in the study were approved by the Institutional Animal Care and Use Committee of the University of Illinois. RGS2 mice breeding cages were taken from Kendal Blumer, Department of Ophthalmology and Visual Science, Washington University School of Medicine and bred in the COMRB of the University of Illinois at Chicago, and C57BL/6 (Jackson Laboratory, Farmington, CT, USA) breeding pairs were bred at the University of Illinois at Chicago. Mice colonies were maintained in a pathogen-free housing facility at the University. Experiments were performed in male and female mice that were between 6-8 weeks old.

#### Induction and Assessment of Lung injury

The lung injury was induced by intratracheal administration of LPS (*i.t.* 0.5 mg/kg) (lyophilized E. coli, Sigma Aldrich # 2880) after anesthetization of the mice with ketamine (100 mg/kg) and xylazine (12 mg/kg). Lung injury was assessed by quantifying extravasation of Evans blue (Sigma-Aldrich # S2129-10G) labeled albumin (Fisher bioreagents # BP9704-100) in lungs versus plasma and lung wet-dry weight ratio as previously (Joshi et al., 2020).

#### Isolation, Culture, and transfection of Bone Marrow-Derived Macrophages

Femur and tibia were flushed with RPMI-1640 media (Gibco), containing 25 ng/ml M-CSF, 10% FBS and 1% antibiotic/anti-mycotic as described previously (Joshi et al., 2020; Rayees et al., 2022; Rayees et al., 2019; Rochford et al., 2021). On 5^th^-day, bone marrow-derived macrophages were incubated in 0.1% FBS containing RPMI for one hour before adding LPS (1.0 µg/ml) with 1% FBS containing media.

BMDM were transfected with vector, RGS2-GFP or Gαq Q209L (UMR cDNA Resource Center, University of Missouri-Rolla # GNA0Q000C0) cDNA using Amaxa Nucleofactor electroporation system (Lonza) as described previously (Mihara et al., 2016; Rayees et al., 2019; Yazbeck et al., 2017). Briefly, BMDM were detached using accutase and spun down with complete media. The indicated cDNA was applied to cells with the transfection reagent and electroporated.

#### Bronchoalveolar Lavage Fluid (BALF) and Alveolar Macrophages collection from mice

Mice were sacrificed after endotoxemia at mentioned time points. BALF was isolated using an 18-gauge blunt needle. Briefly, the sacrificed mice were kept in supine, the trachea was exposed, and 1 mL of cold PBS was slowly injected into the lungs through the trachea and aspirated back; the process was repeated 3-4 times. The collected BALF was centrifuged, and the pellet was dissolved into the RPMI media with 10% serum and plated on plastic dishes. AM were collected 60 min after washing the nonadherent cells with PBS 60 min (Bang et al., 2011). RNA was isolated from the macrophages using a QIAGEN RNA isolation kit (Rayees et al., 2019).

#### RNA sequencing and analysis

Total RNA was extracted using a RNeasy mini kit (Qiagen USA). The purified RNA was further processed to deplete rRNA using RiboMinus^TM^ Eukaryote kit v2 (Thermo Fisher Scientific). RNA libraries were constructed using TruSeq RNA library prep kit v2 (Illumina) and sequenced with Nova seq 6000 for PE150bp. The analysis of raw FASTQ files was carried out as previously described. Briefly, the transcript expression level was calculated and normalized based on counts per million reads (CPM). The differential transcriptome analysis was performed using edgeR with the fold of change ≥1 and false discovery rate (FDR) ≤0.001 considered to be significant differential gene. Volcano plots were plotted between the groups using the Enhanced Volcano R package. Further, gene ontology (GO) biological process functional pathway analysis was carried out using DAVID 2021 at P adjust <0.05 using top 100 upregulated or downregulated genes. The pathway plots were plotted considering the P adjusted to gene count of respective pathways using ggplot2 R package.

#### SARS-CoV-2 virus study

All experiments with live SARS-CoV-2 were performed in a BSL3 laboratory by Zhang et al. (Zhang et al., 2022) after approval by the Office of Environmental Health and Safety at the University of Illinois at Chicago. Briefly, 2019n-CoV/USA WA1/2020 isolate of SARS-CoV-2 (NR-52281) obtained from BEI Resources were propagated in the Vero E6 cells (CRL-1586, American Type Culture Collection, ATCC), and titers were quantitated by a plaque-forming assay (Zhang et al., 2022). Hemizygous K18-hACE mice (strain#034860: B6Cg-Tg(K18-ACE2)2Prlmn/J; Jackson Laboratory), were inoculated intranasally with 1×10^5^ PFU (plaque-forming units) SARS-CoV-2 in 50μL of sterile phosphate-buffered saline. Mouse lungs were perfused and harvested on the 7^th^ day post inoculation for determining RGS2 and IFNγ gene expression.

#### Bone Marrow Transplantation (BMT)

Sub-lethal irradiation (10 Gy) was given to RGS2^-/-^ and WT mice (Weber et al., 2015), followed by transplantation of 2x 10^6^ donor bone marrow cells in 200 µl volume and vice versa (RGS2 null and WT) using a 27-gauge needle. Six weeks after BMT, experiments were performed to assess the lung injury (Rayees et al., 2019; Tauseef et al., 2012).

#### BALF and Alveolar Macrophages collection from ARDS patients and healthy subjects

Healthy subjects and ARDS patients were included in the study. The inclusion criteria for the study were as follows:

1. Age ≥ 18 years, male or female
2. 2. Meet the diagnostic criteria for severe pneumonia (Mandell et al., 2007) or acute respiratory distress syndrome.
3. 3. Provide signed informed consent.

Bronchoscopy and BAL are performed by experienced pulmonologists following a standardized protocol according to current National Heart, Lung, and Blood Institute guidelines (1991). The recovery rate > 30 % (at least 30 mL) is considered a qualified sample. Lavage fluids collected into 50 mL tubes kept on ice and immediately sent to the laboratory by maintaining 4 ^0^C temperature. Next, we filtered the BAL fluid through a 70 μm mesh and centrifuged it at 400 g for 10 min, 4 °C. 500 μL supernatant was used for measuring cytokine. Erythrocytes were lysed using the Red Blood Cell (RBC) lysis buffer. Cells were washed again and resuspended in sorting buffer to process. AM were isolated using an immunomagnetic process. CD14^+^ cells were used as AM with negative selection of EpCAM^-^/CD31^-^/CD66b^-^/CD3^-^/CD20^-^/CD56^-^ cells.

#### Assessment of gene expression by quantitative RT-PCR

To estimate the gene expression following lung injury, total RNA was extracted from the whole lungs or isolated macrophages or BMDM using TRIzol reagent (Invitrogen Inc, Carlsbad, CA). Using Biodrop, RNA was quantified and using specific primers reverse transcription reaction was carried out as per published protocols (Joshi et al., 2020; Rayees et al., 2019).

#### Immunoprecipitation and Immunoblotting

Cells or lungs were lysed in RIPA buffer [150mM NaCl, 10mM-Tris Cl (pH 7.4), 1mM EGTA, 1mM EDTA, 0.5% NP 40, 1.0% Triton X-100, 1mM Na3VO4, 1mM phenyl methyl sulfonyl fluoride (PMSF), 25ml/mg protease inhibitor (Thermo-Scientific, # 1861278). Lysates were incubated with the indicated antibodies overnight at 4 °C. Thereafter, beads were washed with PBS, and cells proteins were separated using Western blotting. Normal rabbit IgG (Santa Cruz # sc-2027) was used for control. The lysates were immunoblotted using indicated antibodies: anti-GFP, anti-phospho-STAT3 (S727 from Santa Cruz # 9134P), anti-phospho-STAT1 (S727 from Santa Cruz # 91775), anti-phospho-STAT2 (Y690 from Santa Cruz # 4441T), anti-phospho-STAT5 (Tyr694, D47E7, from Santa Cruz # 4322), and anti-phospho-NFκB p65 (S536, 9341 from Santa Cruz # 3033S), anti-TLR4 (Santa Cruz #sc-293072), anti-Gαq (Santa Cruz #sc-136181). All antibodies were used at 1:1000 dilutions. Anti-rabbit-IgG-HRP (1:2000 dilution) (Santa Cruz Biotechnology) was used as a secondary antibody. β-actin was used as a loading control. ECL signal was recorded on the ChemiDoc XRS Biorad Imager, and data were analyzed with ImageJ as described previously (Joshi et al., 2020).

#### Gene delivery in mice

RGS2-GFP construct (G14AP26M039) was delivered in mice using cationic liposomes prepared by dissolving Dimethyl Dioctadecyl Ammonium Bromide (MP Biomedicals, LLC # 150939) and cholesterol (Calbiochem #228111) in chloroform, as described (Rayees et al., 2019; Tauseef et al., 2012; Yazbeck et al., 2017). A lipid layer was formed by evaporating chloroform using a rotavapor system (105 rpm for 15-20 min at 37 ^0^C), dissolved in 5% glucose, and extracted by sonicating the solution for 1 h at 37^0^C. Liposomes were filtered using a 0.45-micron filter following which cDNA (vector or RGS2) was added slowly with vortexing to avoid the precipitation of cDNA. The cDNA loaded liposomes was administered retro-orbitally into the mouse for assessment of lung injury.

#### Confocal Imaging

Cells were fixed with 2% paraformaldehyde and incubated with 1% Bovine serum albumin in 0.2% Tween-20 in PBS for 1h at room temperature. After rinsing x3 with PBS, cells were incubated with 1:50 of anti-RGS2 (Santa Cruz # sc-100761), anti-IFNγ-PE (BioLegend # 505807) or anti-TLR4 (Santa Cruz # 293072) antibodies for 1h at room temperature followed by incubation with 1:200 dilution of Alexa-Fluor 594 (Invitrogen # A-21203) secondary Donkey anti-mouse antibody at room temperature for 1h. DAPI (1:1000. Thermo-Fisher Scientific, # D-1306) was used for staining nuclei. The images were acquired with an inverted laser-scanning confocal microscope (LSM 880, Carl Zeiss Microscopy) using the Zeiss LSM software.

#### Cytosolic Ca^2+^ measurement

Cytosolic Ca^2+^ was measured using Fura 2-AM (Invitrogen, # F1221) dye as described (Rayees et al., 2019). Briefly, after maturation, cells cultured on 35 mm glass bottom dishes were incubated with Fura 2-AM dye for 15 mins. Cells were rinsed x2 with HBSS (without Ca^2+^) to deplete intracellular Ca^2+^ after which cells were stimulated with 50 nM thrombin to determine Ca^2+^ influx.

In other studies, BMDM were transfected with GFP-tagged gCaMP-X (Gifted by Dr Borz), which binds Ca^2+^ (Yang et al., 2018). The cells were stimulated with thrombin after 48 h of transfection. The cells were fixed and imaged using confocal microscope.

#### Statistical Analysis

Results are expressed as means ± SD from three independent experiments. Statistical significance was assessed by one-way ANOVA followed by Tukey’s multiple comparisons test and unpaired t-test with Welch’s correction using Graph Pad Prism version 7.0 (Graph Pad Software, La Jolla, CA).

