## Supplementary file for "RGS2 is an innate immune checkpoint for TLR4 and Gαq-mediated IFNγ generation and lung injury"

Supplement Figure 1

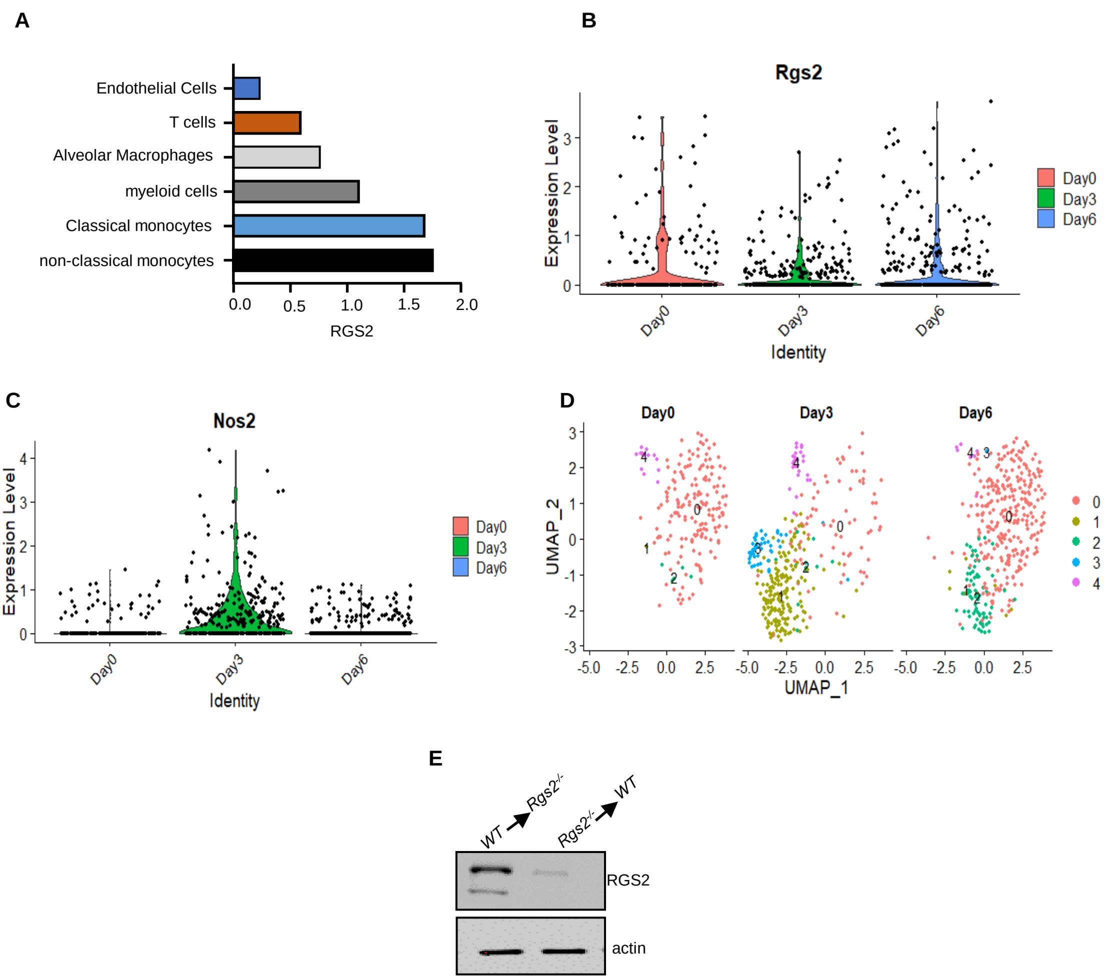

A

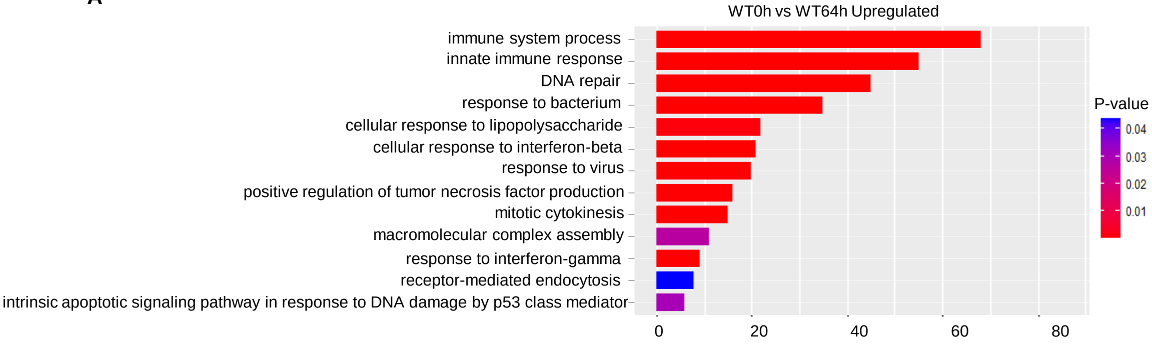
WT0h vs WT64h Upregulated
immune system process
innate immune response
DNA repair
response to bacterium
cellular response to lipopolysaccharide
cellular response to interferon-beta
response to virus
positive regulation of tumor necrosis factor production
mitotic cytokinesis
macromolecular complex assembly
response to interferon-gamma
receptor-mediated endocytosis
intrinsic apoptotic signaling pathway in response to DNA damage by p53 class mediator
0
20
40
60
80
P-value
0.04
0.03
0.02
0.01

B

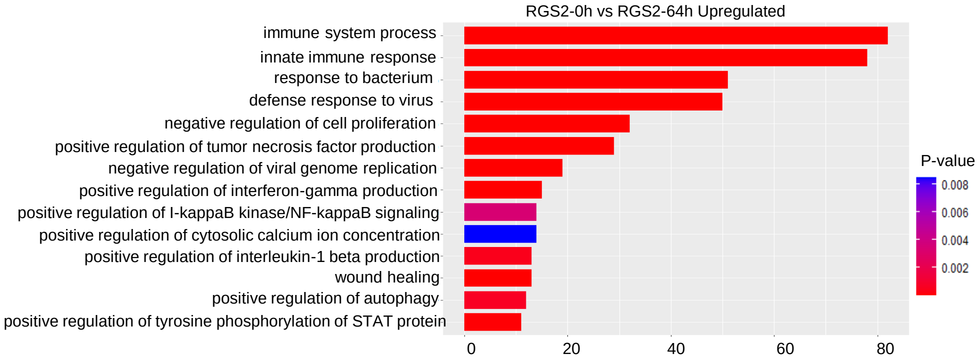
RGS2-0h vs RGS2-64h Upregulated
immune system process
innate immune response
response to bacterium
defense response to virus
negative regulation of cell proliferation
positive regulation of tumor necrosis factor production
negative regulation of viral genome replication
positive regulation of interferon-gamma production
positive regulation of I-kappaB kinase/NF-kappaB signaling
positive regulation of cytosolic calcium ion concentration
positive regulation of interleukin-1 beta production
wound healing
positive regulation of autophagy
positive regulation of tyrosine phosphorylation of STAT protein
0
20
40
60
80
P-value
0.008
0.006
0.004
0.002

C

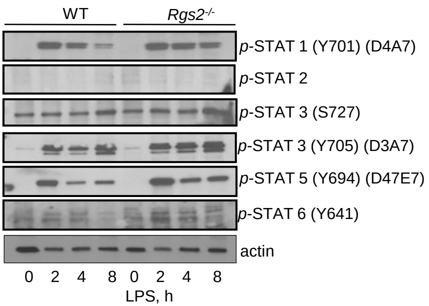
WT
Rgs2-/-
p-STAT 1 (Y701) (D4A7)
p-STAT 2
p-STAT 3 (S727)
p-STAT 3 (Y705) (D3A7)
p-STAT 5 (Y694) (D47E7)
p-STAT 6 (Y641)
actin
0 2 4 8 0 2 4 8
LPS, h

Table 1. Clinical characteristics of 8 ARDS pneumonia patients.

Supplement Figure3

| Characteristics | Pneumonia 1 | Pneumonia 2 | Pneumonia 3 | Pneumonia 4 | Pneumonia 5 | Pneumonia 6 | Pneumonia 7 | Pneumonia 8 |
|---|---|---|---|---|---|---|---|---|
| Age (y) | 73 | 25 | 69 | 20 | 58 | 54 | 86 | 68 |
| Sex | Male | Male | Male | Male | Male | Male | Male | Male |
| Admission date | 08/19/2022 | 08/22/2022 | 08/19/2022 | 08/22/2022 | 07/12/2022 | 08/24/2022 | 09/7/2022 | 09/16/2022 |
| Sampling date | 08/19/2022 | 08/23/2022 | 08/19/2022 | 08/22/2022 | 07/18/2022 | 09/13/2022 | 09/22/2022 | 09/30/2022 |
| Diagnosis | (1) Severe pneumonia (2) Larynx cancer history (Postsurgery) | (1) Community-acquired pneumonia; (2) Hypoxemia. (3) Hypersensitivity pneumonitis (Suspected diagnosis). | (1) Respiratory failure; (2) Pulmonary infections; (3) Pulmonary embolism; (4) Cerebral Infarction; (5) Hypokalemia; (6) Hepatic dysfunction; (7) Cardiac dysfunction; (8) Thrombocytopenia; (9) Urinary tract infection; (10) Diarrhoea; (11) Hypertension; (12) Coronary atherosclerotic heart disease; (13) Amyotrophic lateral sclerosis | (1) Ventilator-associated pneumonia (2) Symptomatic epilepsy (3) Hepatic dysfunction. | (1) Type 2 respiratory failure (2) Interstitial pneumonia (3) Pulmonary embolism (Suspected diagnosis) (4) Hypertension (Grade 3; high risk) (5) Type 2 Diabetes mellitus. | (1) Interstitial lung disease; (2)Dermatomyositis (3)Pneumomediastinum; (4) severe pneumonia; (5) Pleural effusion; (6) Lower extremity venous thromboembolism; (7)Hypokalemia; (8) Hypoproteinemia. | (1) Acute respiratory distress syndrome; (2) Type I Respiratory failure; (3) Spontaneous pneumothorax; (4) Acute interstitial pneumonia; (5) Chronic obstructive pulmonary disease; (6) Severe pneumonia; (7) Pulmonary bulla; (8) Intestinal infection; (9) Grade 3 hypertension (High risk); (10) Hypertensive heart disease; (11) Degenerative heart valve disease; (12) Somatization disorder; (13) Anxiety; (14) Alzheimers disease; (15) Trigeminal neuralgia; (16) Hypoproteinemia; (17) Lower extremity venous thromboembolism | (1) Respiratory failure; (2) Severe pneumonia; (3)Cardiac failure; (4) Major gastrointestinal bleeding; (5) Chronic obstructive pulmonary disease; (6) Asthma; (6) Pulmonary heart disease; (7) Grade 2 hypertension (extremely high risk); (8) Renal insufficiency; (9) Hyperlipidemia; (10) Type 2 Diabetes mellitus. |
| Microbiological diagnosis | Stenotrophomonas maltophilia; Staphylococcus aureus; Klebsiella pneumoniae ; Pseudomonas aeruginosa; Saccharomyces sp. | Undiagnosed | Acinetobacter baumannii ; Pseudomonas aeruginosa; Candida albicans | Klebsiella pneumoniae ;Pseudomonas aeruginosa | Acinetobacter baumannii | Pneumocystis carrinii; Acinetobacter baumannii | Stenotrophomonas maltophilia; Acinetobacter baumannii; pseudomonas aeruginosa; | Acinetobacter baumannii |
| Chest CT Diagnosis | Inflammatory exudation in both lung; Bilateral emphysema. | Scattered inflammatory exudation in both lungs with a few interstitial fibrosis; slightly larger lymph nodes in the mediastinum | Scattered inflammation in both lungs, some with chronic inflammation; atelectasis in the lower lobes of both lungs; and a small amount of pleural effusion on both sides. | Inflammatory exudation in both lung; Lower lobe consolidation in both lungs. | Interstitial pneumonia and fibrosis in both lungs. | Bilateral lung parenchymal exudation; bilateral interstitial inflammation; a small amount of pleural effusion on both sides. | Bilateral emphysema and bulla; cardiac enlargement; minimal pericardial effusion | Inflammation exudation in both lungs, some with chronic inflammation and interstitial fibrosis; Bilateral emphysema; small amount of pleural effusion on the left side |
| Mechanical ventilation | Yes. Invasive | No | Yes. Invasive | Yes. Invasive | Yes. Invasive | Yes. Invasive | Yes. Invasive | Yes. ECMO |
| Oxygenation index | 208.5 | 330.0 | 222.3 | 162.8 | 105.2 | 123.0 | 192.3 | 113.3 |
| Condition (update on | Discharged after improvement | Discharged after improvement | Discharged after improvement | Discharged after improvement | Dead | Discharge after improvement | Under treatment (critically ill) | Dead |

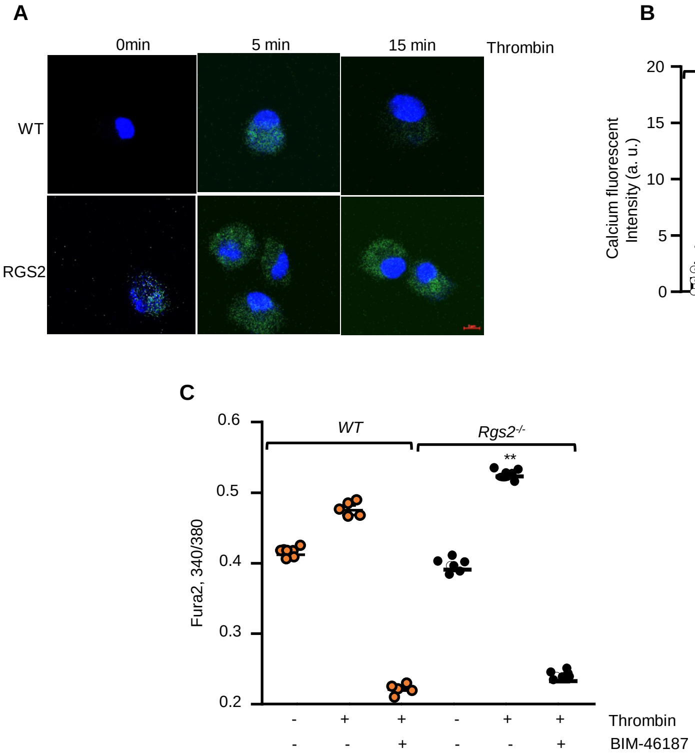
A
0min
5 min
15 min
Thrombin
WT
RGS2
B
WT
Rgs2-/-
**
Calcium fluorescent intensity (a. u.)
20
15
10
5
0
0
5
15
0
5
15
Minutes
C
WT
Rgs2-/-
**
Fura2, 340/380
0.6
0.5
0.4
0.3
0.2
Thrombin
- + + - + +
BIM-46187
- - + - - +

**A**

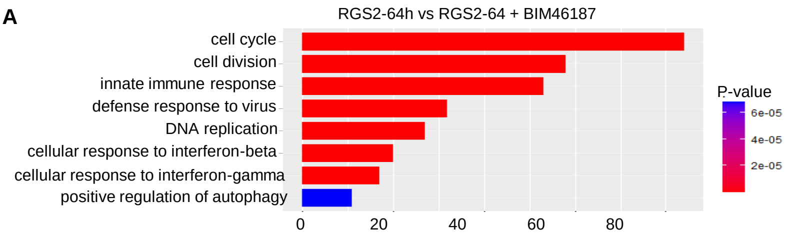

RGS2-64h vs RGS2-64 + BIM46187
cell cycle
cell division
innate immune response
defense response to virus
DNA replication
cellular response to interferon-beta
cellular response to interferon-gamma
positive regulation of autophagy
0
20
40
60
80
P-value
6e-05
4e-05
2e-05


**B**

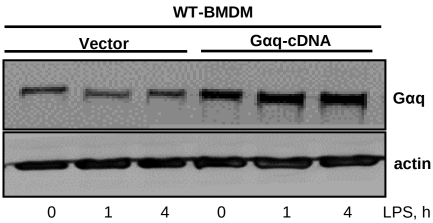

WT-BMDM
Vector
Gαq-cDNA
Gαq
actin
0
1
4
0
1
4
LPS, h
